# Quantifying the vocal repertoire of adult common terns (*Sterna hirundo*)

**DOI:** 10.64898/2026.05.20.722623

**Authors:** Daniel S. Zogby, Valerie M. Eddington, Elizabeth C. Craig, Laura N. Kloepper

**Affiliations:** Department of Biological Sciences, University of New Hampshire, Durham, NH 03824; Center for Acoustics Education and Research, University of New Hampshire, Durham, NH 03824; Shoals Marine Laboratory, joint program of Cornell University Ithaca, NY 14853 and University of New Hampshire, Durham, NH 03824

**Author notes:** Corresponding author: Valerie M. Eddington.

**Keywords:** bioacoustics, acoustic communication, soundscape ecology, seabird communication, signal structure

## Abstract

Common terns (*Sterna hirundo*) are regionally threatened migratory seabirds that form large breeding colonies during the North American summer months. They are highly vocal and serve as important bioindicators of aquatic ecosystems. Historically, acoustic studies on colonial seabirds have proven difficult due to the dense aggregations of individuals and high rate of call overlap. However, as passive acoustic monitoring (PAM) becomes increasingly common for studying seabird colonies, quantitative descriptions of species vocalizations are needed to accurately interpret behavioral information from colony soundscapes and support automated analysis of large acoustic datasets. This study aims to quantify the vocal repertoire of adult common terns. We deployed AudioMoths to collect acoustic data at a tern colony on Seavey Island, New Hampshire, USA from across the breeding season. Using RavenPro, unique call types were identified through visual and aural inspection of the acoustic data in the spectrogram. For each call, we then extracted measurements of peak frequency (Hz), bandwidth 90% (Hz), syllable duration 90% (s), and total bout duration (s) to quantify the characteristics of each call type. Statistical analyses for acoustic parameters by call type were performed using Kruskal-Wallis tests, followed by post-hoc Dunn tests. Our results demonstrate that each call type is significantly different from another by at least one parameter, with the exception of the kek and kip/tjuk calls. These findings present the first quantitative analysis of common tern vocalizations for North America. By defining temporal and spectral characteristics for multiple call types, this work helps translate colony soundscape into biologically meaningful information about tern behavior and colony dynamics. These descriptions also provide key parameters for developing automated tools to detect and classify vocalizations in dense, noisy colonies. Integrating quantified vocal characteristics with PAM offers a promising approach for monitoring colony activity and behavior while minimizing disturbance relative to traditional methods.

---

For many species, acoustic communication plays a critical role in regulating biological interactions, including mating, parent-offspring communication, territorial defense, and antipredator behavior. These functions are especially pronounced in social species such as colonial seabirds, which spend extended periods in large, dense breeding colonies with hundreds to thousands of conspecifics (Nelson and Baird 2001, Schreiber and Burger 2002). When in breeding colonies, seabirds use a combination of vocalizations and postures to convey motivation to nearby conspecifics and perceived threats (e.g., human researchers, predators, Nelson and Baird 2001). Communicative behavior in colonial seabirds can be broadly categorized as either ritualized or overt. Ritualized behaviors are stereotyped, exaggerated, and repetitive, and include behaviors associated with nest acquisition and maintenance, pair formation and bonding, pre- and post-copulatory displays, and parent-offspring communication (Beer 1980, Nelson and Baird 2001). In contrast, overt behaviors such as fighting, fleeing, and feeding have more transparent motivation and function (Nelson and Baird 2001). Regardless of behavioral category, the diverse vocal behaviors of colonial seabirds play a crucial role in colony fitness and are a defining feature of seabird breeding colonies.

The common tern (*Sterna hirundo*) is a globally distributed migratory seabird. Here, we focus on a costal-breeding population in the Northwest Atlantic that breed in North America and winter along the coasts of Central and South America (Neves et al. 2015, Caldwell et al. 2025). Like other colonial seabirds, common terns exhibit a diverse and context-dependent vocal repertoire that facilitates mating, territory defense, chick provisioning, and predator deterrence (Arnold et al. 2020).

Common terns produce different vocalization types associated with behavioral state, and these call types have been phonetically described in the literature. The “kee-ar” or “kyar” call is typically associated with escape and alarm behavior, performed most often when terns are flying overhead the colony (Veen 1987; Arnold et al. 2020). The “long call,” also called the “advertisement call,” is a general-purpose call noted to vary among individuals, potentially used for individual recognition (Stevenson et al. 1970), and is often associated with the return of a parent to the nest site, provoking chicks to emerge from hiding places (Veen 1987; Arnold et al. 2020). “Kip” or “Tjuk” calls are thought to express low-level aggression with conspecifics, produced most often when individual’s take-off from the ground (Veen, 1987; Arnold et al. 2020). Fear calls, described as a descending “kyew,” are produced when startled by an intruder or during aggressive conspecific encounters. Brooding calls are lower in frequency, described as a crooning call “krr-krr-krr-krr” and produced when tending to the nest, eggs or chicks (Arnold et al., 2020). “Kek,” or attack calls, described as rattle and growl call (Veen 1987) with a “kek-kek-kek-kek” structure, are given during attacks of humans or conspecifics, often ending in a “kyar” call during intruder strike (Arnold et al., 2020). “Korr-korr-korr” calls are lower in frequency than most calls and emitted during both high flights and when an intruder approaches an adult at the nest. Copulation calls are described as “quiet, squeaky kyi-kyi-kyi-kyi” and emitted by males during both precopulatory displays and when engaged in mounting (Arnold et al., 2020).

Begging calls, described as “ki-ki-ki-ki-ki-ki” are produced by chicks when soliciting food from parents, but these calls are also made by adults of both sexes during close interactions at the nest, leading to the theory that these vocalizations indicate appeasement (Arnold et al., 2020). Chicks are known to produce a “peep” at early stages of development, (∼24 hours pre-hatch to 2-3 days post-hatch), transitioning to a “cheep” (3-6 days post-hatch), followed by “distress” and “begging” calls that continue for 2-3 months post-fledging (Arnold et al. 2020).

Despite the well-documented behavioral context, the existing literature on common tern vocalizations consists largely of qualitative descriptions or limited quantitative analyses of call structure (Tinbergen 1931, Stevenson et al. 1970, Glutz von Blotzheim and Bauer 1982, Becker and Ludwigs 2004, Arnold et al. 2020). While these studies provide valuable insights into general communication behavior, new technologies now allow the vocal repertoire of the common tern to be more quantitatively characterized using standardized acoustic parameters commonly applied in modern acoustic ecology (e.g., Gleason et al. 2023, Eddington et al. 2024, Zager et al. 2024).

A comprehensive quantitative description of vocalizations across call types can facilitate standardized acoustic measurement, improve comparability across studies, and support the development of automated detection tools. Such characterization is particularly important for advancing passive acoustic monitoring (PAM) applications in this species. PAM offers a minimally-invasive approach to survey vocal species and enable the long-term monitoring while reducing disturbance associated with human presence (Sugai et al. 2019, Brosseau et al. 2024), an important consideration for seabird colonies where disturbance can negatively affect reproductive success {Citation}(Carney and Sydeman 1999, Blackmer et al. 2004, Shonfield and Bayne 2017). Colonial seabirds are particularly well suited for PAM due to their high vocal activity rates, and PAM has been successfully applied to estimate population density and characterize behavioral patterns in other seabird species (Borker et al. 2014, Descamps and Ramírez 2021, Patterson et al. 2022). Furthermore, quantitative call descriptions are necessary for the development of automated detection and classification tools (e.g., BirdNET; Kahl et al. 2021) and for colony-level soundscape analyses using acoustic indices (Bradfer‐Lawrence et al. 2024).

The goal of this study is to describe the quantitative characteristics of the adult vocalizations of common terns within a northwest Atlantic breeding colony. Specifically, we aim to identify and describe distinct call types based on measurable acoustic parameters and match each unique call type with the existing qualitative descriptions in the literature. We hypothesize that adult call types can be distinguished quantitatively and predict that each identified call type will differ significantly from other call types by at least one parameter. By doing so, we provide the first quantitative vocal repertoire for the common tern, laying the groundwork for future behavioral, ecological, and passive acoustic monitoring applications in this species.

## Methods

### Study Area

This project took place across five sites on Seavey Island, NH in the Isles of Shoals (Figure 1). The site, which encompasses both Seavey and White Islands, is a crucial breeding ground for common terns and at approximately 3,100 breeding pairs is the largest active breeding colony for this species within the Gulf of Maine. The Seavey Island colony is a mixed-species seabird colony; common terns are the most populous species, with a smaller nesting population of roseate terns (*S. dougallii;* ∼150 breeding pairs). Small populations of common eiders (*Somateria mollissima)*, spotted sandpipers *(Actitis macularius)*, song sparrows *(Melospiza melodia)*, and arctic terns (*S. paradisaea*) have been observed nesting on Seavey Island in recent years. Herring gulls *(Larus smithsonianus)* and great blacked-backed gulls *(L. marinus)* have been noted on the island as well, although as a result of active management, no gulls nest at this site.

**Figure 1.**
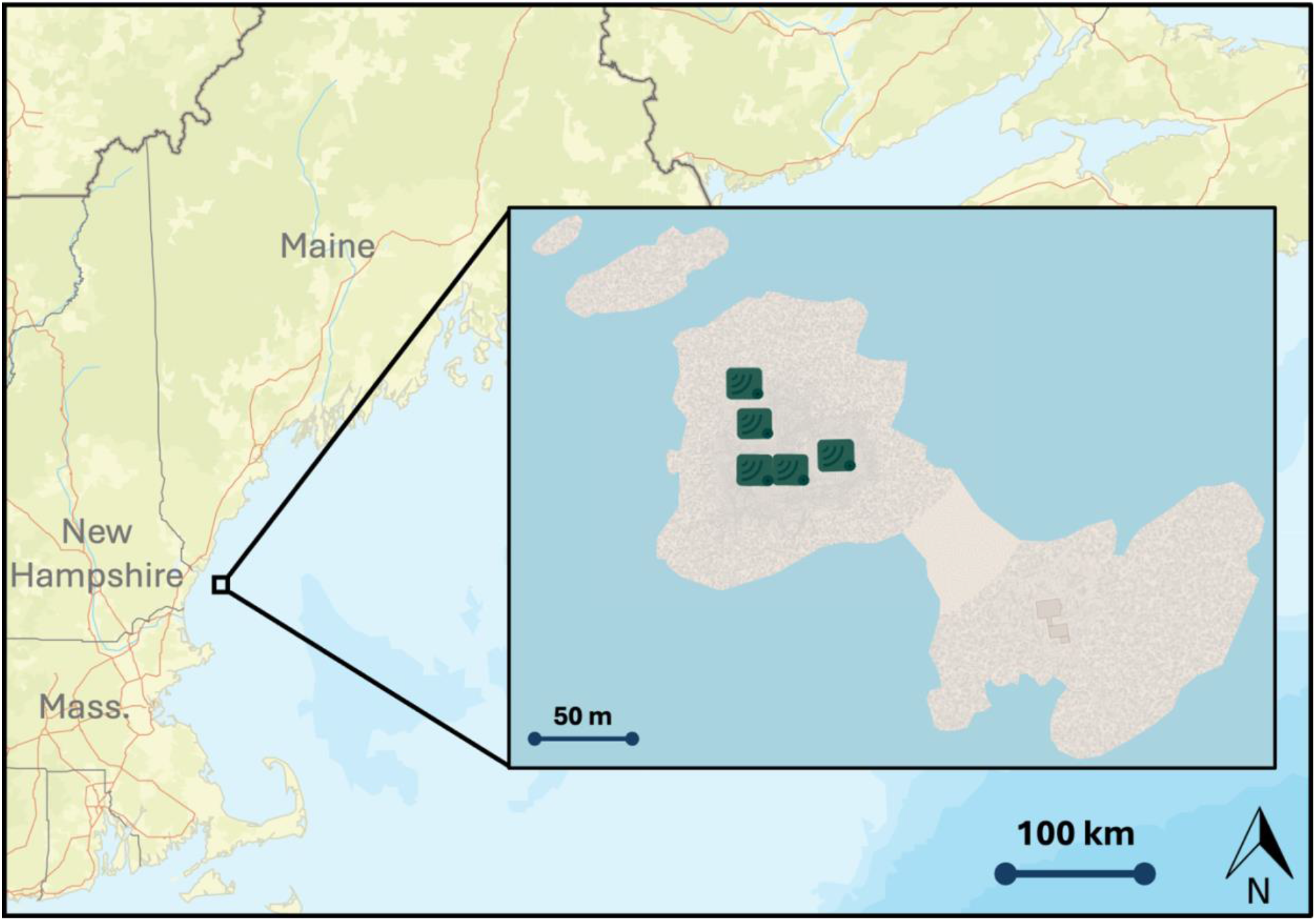
The inset map shows Seavey Island (left) and White Island (right), New Hampshire, located within the Gulf of Maine approximately 6 miles off the coast of mainland New Hampshire, USA. White and Seavey Island are connected by a land bridge at low tide. Green, rectangular icons on Seavey Island represent location of AudioMoth acoustic recording devices in this study.

### Data Collection

We deployed five AudioMoths across the five sites on Seavey Island, NH, recording from 15 May to 7 August 2023. Recorders were attached to PVC pipes and staked into the ground to hold recorders approximately 30 cm above the ground. We programmed Audiomoths to record the first 10 minutes of every hour, 24 hours a day (sampling rate = 48 kHz, low gain sensitivity). For analysis, we subsampled one 10-minute file per day at 1300 h (daytime) EST and one at 0200 h (nighttime) within the same 24-hour period. We screened selected files to ensure the absence of wind or rain noise. If either the daytime or nighttime file for a given 24-hour period contained noise contamination, both files from that day were excluded to avoid potential bias. We targeted three paired day-night samples per week; however, due to weather-related exclusions, some weeks yielded only two usable pairs for final analysis.

### Call Identification

We used RavenPro Acoustic Analysis Software (v. 1.6.5, Cornell Lab of Ornithology, Ithaca, NY) to generate spectrograms from our recordings. Spectrograms were visually and aurally inspected to identify vocalization samples for analysis. We grouped identified vocalizations into two main categories – syllables and bouts. A syllable is a single vocalization, and a bout is a group of consecutive syllables of one call type produced by an individual tern (Figure 2).

**Figure 2.**
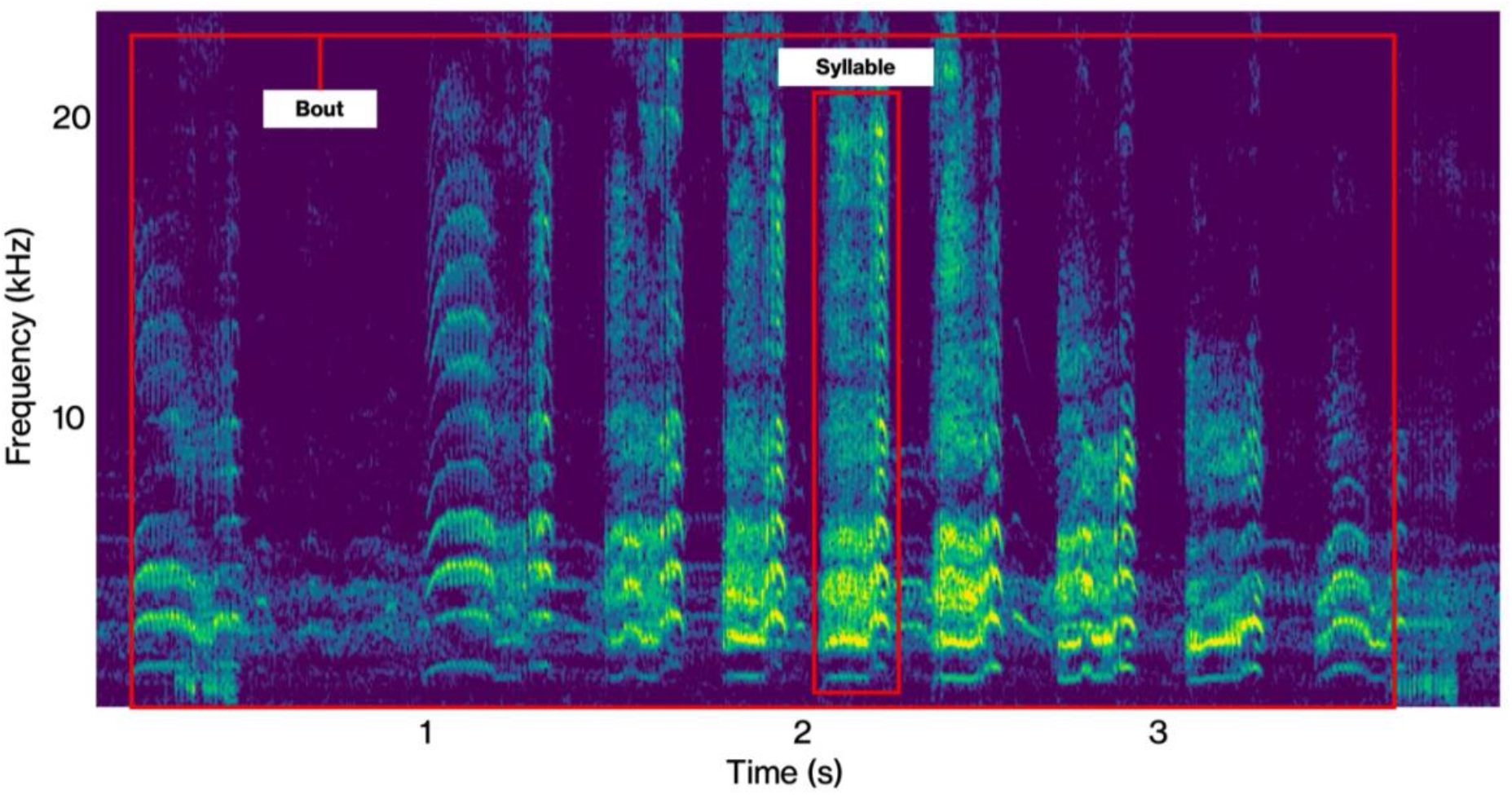
Visual depiction of the distinction between a syllable and a bout (See Methods, *Call Identification*). Spectrographic data displays an example of the long call of a common tern.

We identified high-quality vocalizations by isolating the signal of interest from overlapping vocal activity and calculating the signal-to-noise ratio (SNR). For each bout, the syllable with the highest SNR was selected for measurements of peak frequency, bandwidth, and syllable duration. This approach minimizes measurement error resulting from frequency-dependent attenuation of signal components, which disproportionately affects low-amplitude vocalizations presumed to originate from greater source distances. For spectral analyses, we only included bouts in which the highest SNR syllable exceeded a SNR threshold of 15 dB. Vocalizations falling below this threshold were nevertheless included in bout length analyses, as this temporal metric is not susceptible to frequency-dependent attenuation artifacts. We screened all syllables for audio clipping, a form of signal distortion arising when vocalizations produced at close proximity exceed the dynamic range of the recording microphone. Consequently, we excluded all syllables containing clipping artifacts from all subsequent analyses.

For each vocalization, we extracted the total bout duration. For the highest SNR syllable within each bout, we extracted peak frequency (kHz), bandwidth (kHz), and syllable duration (s). We measured these acoustic parameters using the 90% signal energy criteria built into Raven Pro. We then categorized bouts into the phonetic descriptions of common tern vocalizations as described in the literature (Cramp 1985, Veen 1987, Becker and Ludwigs 2004, Arnold et al. 2020). Because chick vocalizations are typically quieter than adults (Arnold et al. 2020), we focused our efforts on adult vocalizations and did not attempt to classify or analyze chick calls (i.e., calls we identified as “peep” and “cheep”).

Because visual observations were not paired with acoustic recordings, we based species identification on auditory characteristics of calls. Prior to analysis, the observer spent time in the colony becoming familiar with the vocal repertoire of all species present on the island, including the co-nesting roseate tern, and reference recordings from the MacCaulay Library were used to confirm species association when needed. Although most vocalizations between common and roseate terns are readily distinguishable, some call types are more acoustically similar than others, particularly the kek call of the common tern and the attack call of the roseate tern, which share a similar phonetic description (Arnold et al. 2020, Gochfeld and Burger 2020). We therefore classified calls conservatively and excluded uncertain vocalizations from analysis.

### Statistical Analyses

We conducted all statistical analyses and data visualization in RStudio (v. 2025.05.1, Posit Software, Austria). To assess statistical differences in the acoustic parameters of peak frequency (kHz), bandwidth 90% (kHz), and duration 90% (s) for both syllables and bouts, we used Kruskal-Wallis tests, supplemented by a post-hoc Dunn test. For syllable analysis, we filtered bouts to select only the syllable with the highest SNR for peak frequency, bandwidth, and syllable duration analysis to prevent pseudo-replication. We included all syllables in the analysis of bout length. For syllable duration, peak frequency, and bandwidth, a Bonferroni correction was applied to the α-value (α = 0.0167) used to assess p-value significance to control for family-wise errors for analyses sharing the same dataset. For total bout duration, we used an α = 0.05 because the dataset used for this analysis differed from those used for the other acoustic parameters.

## Results

We identified six categories of vocalizations that we matched to the phonetic descriptions in the literature: kee-ar, long call, kip/tjuk, fear call, brooding call, and kek. Representative spectrograms for each call type category are shown in Figure 3, with summary characteristics of each call type listed in Table 1. On average, bout durations ranged between 0.4 and 4 seconds, syllable durations ranged between 0.01 and 0.7 seconds, peak frequencies ranged between 3 to 5 kHz, and bandwidths ranged between 3.8 and 12 kHz.

**Figure 3.**
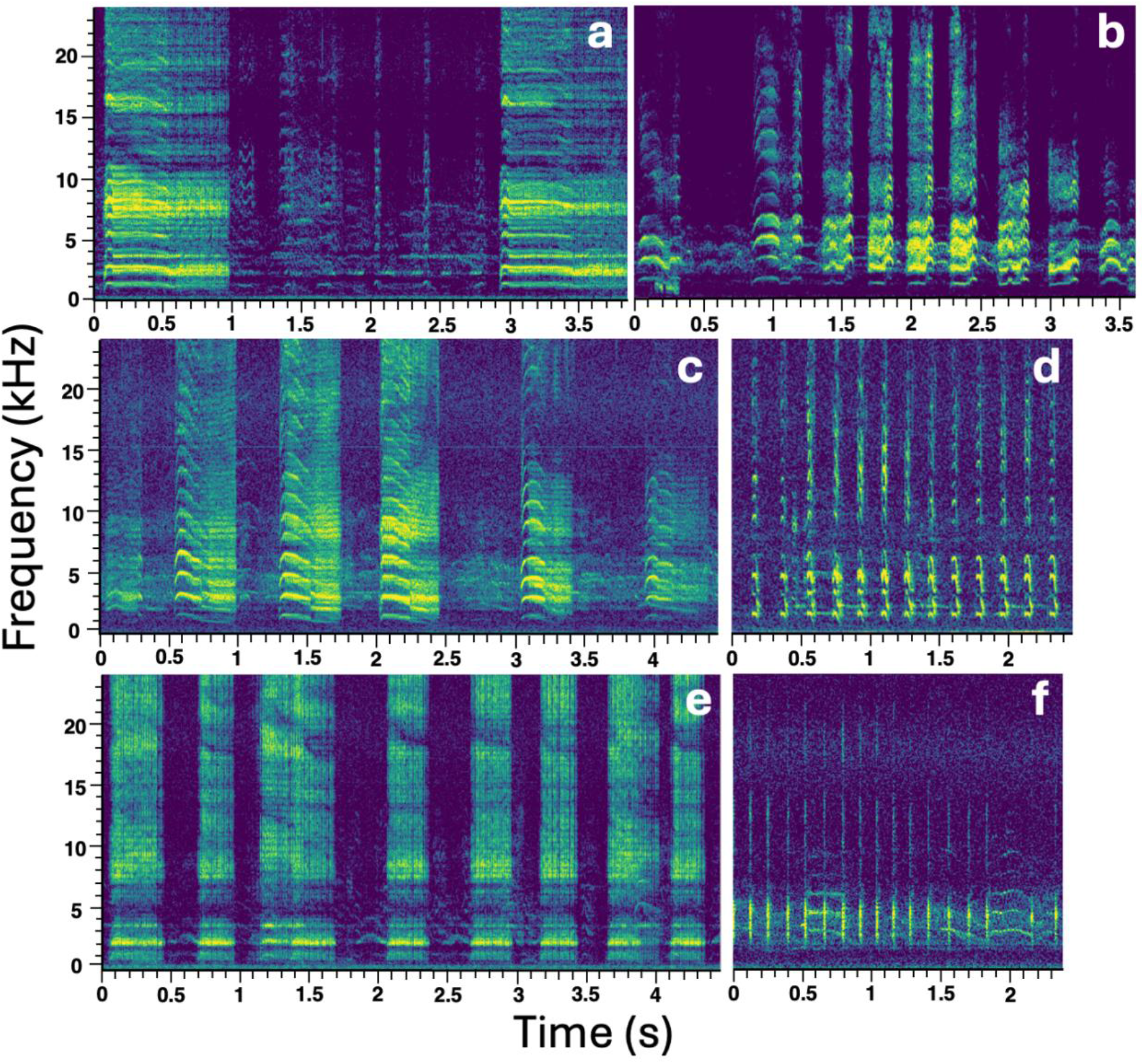
Representative spectrograms of identified unique common tern vocalizations from our dataset. (a) kee-ar, (b) long call; (c) fear call, (d) kip, (e) brooding call, and (f) kek.

**Table 1:** Number of vocalizations (N) along with median (Interquartile range) of measured acoustic parameters associated with each call type.

| Call Type | N | Bout duration<br>(s) | Syllable<br>duration (s) | Peak frequency<br>(kHz) | Bandwidth<br>(kHz) |
| --- | --- | --- | --- | --- | --- |
| Kee-ar | 31 | 0.83 (0.34) | 0.69 (0.27) | 4.59 (2.58) | 7.97 (9.66) |
| Long Call | 117 | 1.90 (2.76) | 0.27 (0.09) | 3.89 (1.97) | 6.42 (4.85) |
| Fear Call | 100 | 0.44 (0.94) | 0.28 (0.08) | 4.55 (1.92) | 6.66 (6.52) |
| Kip/Tjuk | 133 | 1.17 (2.00) | 0.03 (0.01) | 4.97 (1.69) | 8.44 (8.72) |
| Brooding Call | 20 | 3.99 (5.02) | 0.13 (0.12) | 3.28 (1.73) | 3.80 (9.54) |
| Kek | 15 | 1.00 (1.66) | 0.01 (0.01) | 4.69 (2.06) | 12.00 (7.92) |

A Kruskal-Wallis test for each parameter found significant variation across call types for bout duration (s) [X^2^ = 44.618, p < 0.01], syllable duration 90% (s) [X^2^ = 335.79, p = <0.01], peak frequency (Hz) [X^2^ = 28.148, p = < 0.01], and bandwidth 90% (Hz) [X^2^ = 15.554, p = < 0.01] (Figure 4). Post-hoc Dunn tests were used to evaluate pairwise differences among call types. While the Kruskall-Wallis test indicated significant differences in bandwidth among call types (p<0.01), the post-hoc Dunn test with Bonferroni correction did not identify any pairwise differences that remained significant after correction (Figure 4D). Overall, each call type differed significantly from at least one other call type in at least one of the parameters (p<α), with the exception of the kip/tjuk and kek calls which were not significantly different across any of the four parameters evaluated (p>α; Figure 4).

**Figure 4.**
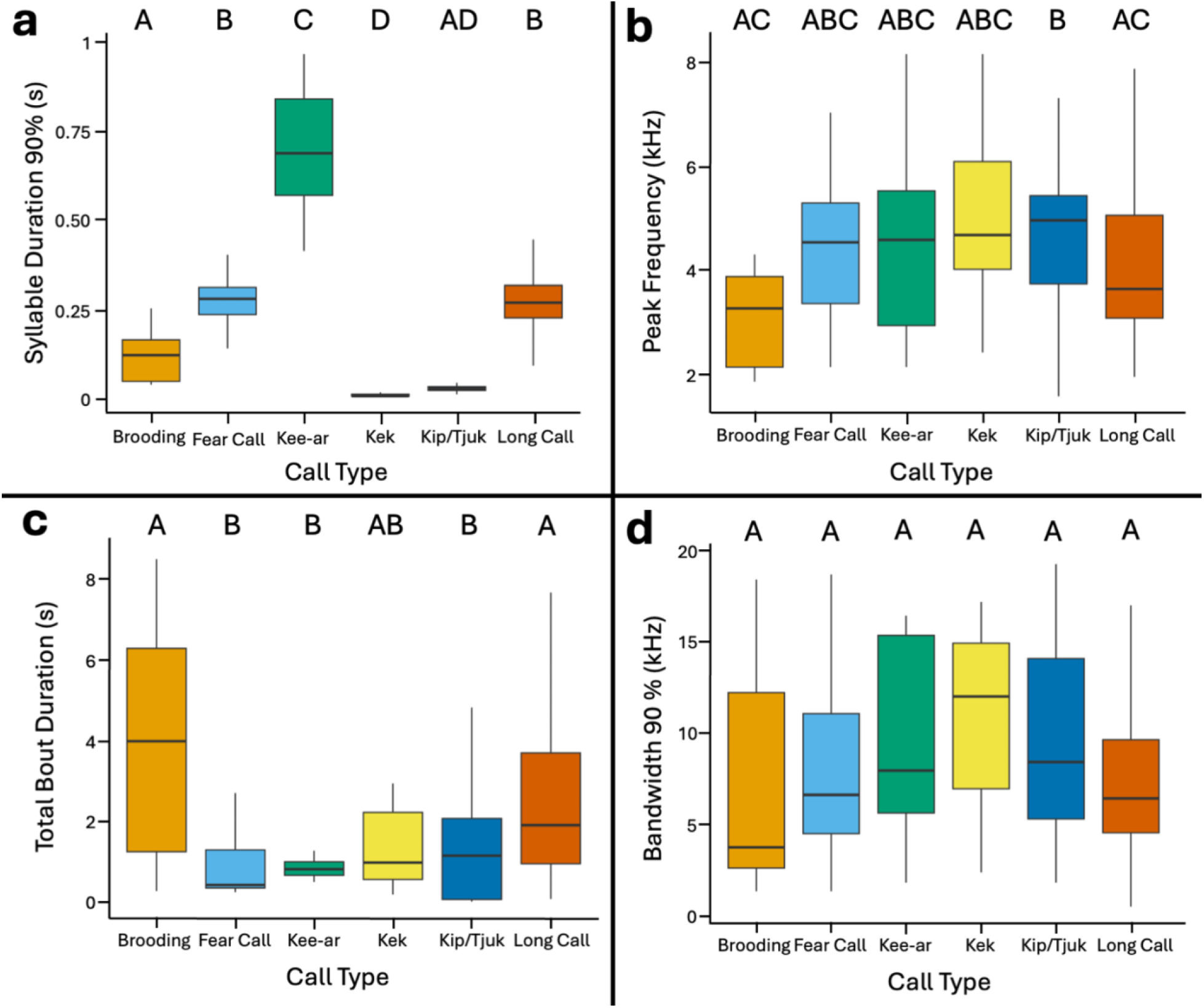
Variation in syllable duration 90% (s) (a), peak frequency (kHz) (b), total bout duration (s) (c) and bandwidth 90% (kHz) (d) across call types. For all panels, bold horizontal lines represent median values, and the surrounding box shows the interquartile range. Vertical lines represent first and fourth quartiles. Outliers were removed to preserve scaling. Letters represent groups statistically significant from each other according to Dunn test; for all panels except total bout duration (c), significance is based on a Bonferroni-adjusted α = 0.0167, while for total bout duration, significance is based on α = 0.05.

## Discussion

Our findings support our hypothesis, with each adult common tern call type varying significantly from other call types by at least one parameter, with the exception of the kek and kip calls. Comparing the different call types we detected, kee-ar calls have some of the shortest bout durations but longest syllable durations, whereas brooding calls demonstrate the opposite trend, with longer bout durations but shorter syllables. The values we report for bouts and syllables (Table 1) provide useful time and frequency ranges for developing automated classifiers of tern vocalizations, and the summary parameters for each call type offer quantifiable metrics for calls that are often described only phonetically in the literature. It is important to note that we did not detect all adult vocalizations described in the literature. Specifically, korr-korr and copulation calls were absent from our dataset. Given the similarity of phonetic descriptions between korr-korr and brooding calls, it is possible these call types were grouped together in our analysis. Similarly, copulation calls may have gone undetected because they are described as “quiet” vocalizations (Arnold et al., 2020) and may have been masked by background colony noise.

Peak frequencies and bandwidths experienced relatively large variation across call types relative to their mean (Figure 4). Such variation is expected because seabird vocalizations serve diverse functions, including individual recognition, parent-offspring communication, mating displays, territory defense, and antagonistic/antipredator displays (Stevenson et al. 1970, Palestis and Burger 2001, Arnold et al. 2020). Both individual-variation and context-dependent variation would lead to diversity in acoustic parameters within a given call type. Flexibility in call structure may also allow individuals to adjust signal characteristics (e.g., frequency or amplitude) to environmental conditions such as temperature, habitat structure, or background noise in order to maximalize signal propagation and detectability (Nottebohm 2013). In dense and acoustically complex breeding colonies, modifying spectral or temporal features of calls could increase the probability that signals reach their intended receivers.

For calls involved in individual recognition, such as the long call (Stevenson et al. 1970, Cramp 1985, Arnold et al. 2020), high variability in acoustic parameters is expected because distinctive features facilitate discrimination among individuals (Carlson et al. 2020). In socially monogamous colonial seabirds, such variability across several call types may help maintain strong pair bonds and social relationships among neighbors across breeding seasons (Osiecka et al. 2024). However, flexibility in acoustic structure is likely advantageous across call types, beyond just individual recognition. Alarm signals, for example, may exhibit graded variation in intensity (e.g., call rate or amplitude) to convey information about threat level and to elicit appropriate responses from nearby conspecifics (Suzuki 2014). The kee-ar and kek calls, both associated with escape and alarm behavior in common terns, may function as graded alarm calls and therefore benefit from flexibility in structure and/or intensity (Veen 1987; Arnold et al. 2020). One study documented increased common tern call amplitude in response to human presence within a colony, although it remains unclear whether this pattern reflected increased calling rate, more individuals calling, or higher amplitude of individual calls (Brosseau et al. 2024). Finally, calls associated with conflict and territorial interactions, such as the kip/tjuk call (Arnold et al. 2020), may also vary in acoustic parameters if such variation communicates information about individual condition or competitive ability, potentially influencing outcomes of aggressive encounters and mediating conflict resolution (Bradbury and Vehrencamp 2011).

The lack of clear differentiation between kek and kip/tjuk calls across the measured acoustic parameters may reflect functional overlap between these vocalizations. Both calls occur in high-arousal contexts associated with aggressive or territorial encounters with conspecifics or predators (Veen 1987, Arnold et al. 2020), and their acoustic structure may prioritize signal flexibility over strong differentiation in temporal and spectral parameters quantified here. However, aural and visual inspection of the calls indicate that these calls differ in spectral structure in ways not captured by our measured parameters. Although the two signals share similar peak frequency, duration, and bandwidth, kip calls exhibit a clear harmonic structure (Figure 3C), whereas kek calls appear more homogenous across frequencies, lacking harmonic components (Figure 3F). These differences may indicate distinct sound-production mechanisms. In fact, while kip calls are likely produced through typical syrinx-based vocalizations, we theorize the kek call is produced by clacking the upper and lower halves of their bill together very quickly, rather than coming from the syrinx. Because this study relied solely on acoustic recordings, we could not confirm the behavioral mechanism producing the kek call. Future work pairing synchronized video and acoustic recordings would help clarify the production mechanism and functional differences between these signals.

It is important to acknowledge a few limitations of this study. First, we were analyzing birds during their breeding season at a single location on the ground, so we were almost certainly not comprehensive in identifying every call type produced by adult common terns. Given the vocal activity of the breeding colony, calls produced by birds in flight were not likely to be received at SNR values high enough to be detected by our recorders. Furthermore, it is reasonable to assume that terns may make different vocalizations outside of the breeding colony, especially during periods of feeding or contact at sea. Second, this study relied solely on acoustic data. The absence of visual observations to complement our acoustic data made it impossible to determine the individual producing the vocalizations and to explicitly link specific behaviors to call types. Future studies should pair acoustic data with visual components (e.g., blind observations, camera traps) in order to provide explicit linkages between vocalizations and corresponding behaviors. Finally, we did not attempt to quantify or categorize chick vocalizations. Future work could investigate ontogenetic changes in vocalizations and the role of chick and juvenile vocalizations in parent-offspring and conspecific communication.

Despite these limitations, this quantification of tern vocalizations provides an important foundation for advancing PAM of common tern breeding colonies. By establishing temporal and spectral ranges for multiple call types, this work helps translate colony soundscape into biologically meaningful information about tern behavior and colony dynamics. These quantitative descriptions also provide critical parameters for developing automated detection and classification tools capable of identifying specific vocalizations within dense, noisy colonies. As acoustic monitoring becomes increasingly common in ecological research, linking soundscape metrics to well-described species vocal repertoires will be essential for interpreting long-term acoustic datasets. For colonial seabirds such as common terns, integrating quantified vocal characteristics with PAM approaches offers a promising pathway for monitoring colony activity, phenology, and population trends while minimizing disturbance compared to traditional survey methods.

## Acknowledgments

We would like to thank Aliya Caldwell, Willow Dalehite, Orena Wong, and Joseph Brosseau for assistance deploying and recovering AudioMoths and collecting field data. We would like to thank all current and previous members of the Ecological Acoustics and Behavior Lab for their continued support. This manuscript is contribution #217 to the Shoals Marine Laboratory.

## Conflict of Interest

The authors have no conflicts to disclose.

## Funding

This project is funded by a National Science Foundation Award 2226886, the UNH School of Marine Science and Ocean Engineering Graduate Student Research Fund, the UNH Collaborative Research Excellence Initiative, and the UNH Hamel Center for Undergraduate Research. The Isles of Shoals Tern Conservation Program is supported by the New Hampshire Fish and Game Department’s Nongame and Endangered Wildlife Program, the United States Fish and Wildlife Service (New Hampshire State Wildlife Grants), and the Shoals Marine Laboratory.

## Permits

The authors have no conflicts to disclose. Tern colony fieldwork was permitted under protocol #220304 approved by University of New Hampshire’s Institutional Animal Care and Use Committee.

## Statement on the Use of Generative AI

During the preparation of this manuscript, the authors used generative AI tools to assist in copy-editing of drafted text and troubleshooting code for data cleaning. For copy-editing assistance, the authors prompted ChatGPT versions 5.1-2 with drafted sections, requesting specific improvements to grammar and clarity without altering scientific content. The model returned revised text, which the authors reviewed and selectively incorporated, verifying that meaning, terminology, and citations remained unchanged. For data cleaning, the authors prompted ChatGPT versions 4.1, 5.1-2 with their data cleaning code and associated observed issues and returned error messages. The model returned suggested code modifications, which the authors selectively implemented. The authors verified these changes by manually validating code behavior on subsets of the data and comparing code outputs to expected values to ensure that data cleaning procedures were implemented correctly and produced the intended results. The authors maintained full responsibility for the content, interpretation, and accuracy of all text and analyses.

## Notes

### Competing Interest Statement

The authors have declared no competing interest.

